# The genetic and environmental basis of hematopoietic chimerism in marmosets

**DOI:** 10.64898/2026.11.19.767351

**Authors:** Ian H. Cheeseman, Alexana J. Hickmott, Catherine Jett, Martha Lyke, Matthew Lopez, Alec Bagwell, Riti Sharan, Kimberley A. Phillips, Kelly R. Reveles, Angélica Olmo-Fontánez, Jessica Grieg, Deborah Newman, Donna Layne-Colon, Julienne N. Rutherford, Shelley Cole, Corinna N. Ross

## Abstract

Common marmosets (*Callithrix jacchus*) have a high rate of twinning during pregnancy and exhibit an unusual phenomenon whereby stem cells migrate between fetuses *in utero*. These cells are integrated into a growing sibling resulting in each individual being a chimera of self and sibling. We mapped the distribution of chimerism across 145 marmosets by amplicon sequencing, showing high levels of chimerism in lateral mesoderm-derived tissues. By integrating pedigree and clinical records with estimates of chimerism we identified litter size to be a major determinant of chimerism and lifespan and demonstrate that chimerism and litter size are both heritable traits. Using exome sequencing of ∼400 marmosets we identified major loci associated with variation in litter size and chimerism. These findings have implications for our understanding of self- and non-self across evolutionary biology, immunity and medicine.

---

Chimerism occurs when an individual is composed of cells, and thus genetic material, of multiple individuals^1^. In mammals, chimeric transfer of genetic material can occur via transplacental sharing between mother and offspring, via dizygotic twin-twin transfusion, inheritance via the germline, or artificially following transfusion or organ transplant^2,3^. With the availability of affordable genomic tools, there is an increase in the known instances of chimerism in humans. One study found chimerism in 8% of 415 twin pairs and 21% of 57 triplet sets^4^, increasing to ∼90% for monochorionic twins who share a placenta^5^. In 2022, nearly 43,000 people in the United States received organ transplants, and in 2017, over 10.5 million whole blood-derived and apheresis RBC units were transfused providing an abundant opportunity for chimerism to arise^6^.

Chimerism has long been understood to be a widespread phenomenon in marmoset monkeys. Female marmosets release oocytes from both ovaries asynchronously^7^. This produces multizygotic litters, typically twins in the wild, but triplets, quadruplets and even quintuplets are common in captivity^8^. Regardless of litter size, the marmoset placenta comprises two discs connected via blood vessels; marmosets, unlike other mammals, are thus monochorionic and multizygotic. This unusual combination means that multizygotic littermate-littermate transfusion, is a physiologic feature of marmoset development and as a result, chimerism is highly prevalent.

Chimerism in marmosets has been assessed by karyotyping, microsatellite genotyping, and Y chromosome genotyping. There is a consensus that ∼90% of marmoset pregnancies result in chimerism of hematopoietic tissues (e.g. placenta, blood, spleen, bone marrow) in offspring^9–13^, but a persistent question is whether other tissues are also chimeric, and to what degree. The presence of sibling cells appears to be idiosyncratic among tissues, individuals, and litters, and the causes of this variation and the timing and sequence of cell transfer remain unknown. It was initially thought that primordial germline cells migrating from the yolk sac to the genital ridges could detach and enter shared fetal circulation though no evidence was initially found for germline chimerism^11^. Using DNA fingerprinting Signer *et al*. were unable to detect chimerism in non-hematopoietic tissues, though the study was limited by low resolution techniques^13^. Subsequent analysis of tissues for the SRY gene concluded that germ line cells were not chimeric and only hematopoietic lineages were^14^. Conversely, a novel evaluation of microsatellites from pregnancies with known parentage and samples from multiple tissue lineages estimated that all tissues were potentially chimeric, including germ line cells in nearly 60% of animals tested^12^. Even if only hematopoietic tissue is chimeric in marmosets, these cells interact with all other cells in the body and are responsible for cell-cell communication, delivery of nutrients, immune function, and overall homeostasis, which might have significant influence on tissue function in these animals.

While the impact of chimerism on the function of tissues and health outcomes remains underexplored, there is mounting support for a role for chimerism impacting human health^1,15,16^. Genomic conflict within the body due to microchimerism can trigger an inflammatory cascade and is a risk factor for rheumatoid arthritis,, Grave’s, and Hashimoto’s diseases in women^1,15,17^. Autoimmune diseases have a significant sex bias with some diseases reported as more than 70% occurrence in women, and predominantly in women following childbearing age^18^, likely due to microchimerism^1^. Pathologies associated with transplantation and other sources of chimerism result from chronic graft-versus host disease^2^. Conversely, there are reduced risks for cancer, Alzheimer’s, multiple sclerosis, malaria and respiratory pathogens^16,19–21^ associated with protective functionality of the non-self-cells. In marmosets chimerism has been hypothesized to be the underlying cause of chronic lymphocytic enteritis^22^. The impact of increased non-self-cell contribution due to higher order litters (triplets and greater) remains unclear.

In this study, we present a genomic investigation of the genetic basis of chimerism in the common marmoset. We address (i) the extent and tissue specificity of chimerism, (ii) the factors which influence the presence of chimerism in an individual, and (iii) the host genetic determinants that influence engraftment and persistence of sibling-derived hematopoietic stem cells. By investigating the genetic architecture of hematopoietic chimerism in a primate, we aim to provide insights into both fundamental developmental biology and translational models relevant to human health.

## Results

### Assessment of chimerism in a marmoset colony

We performed genotype-by-sequencing on hair follicle DNA from 82 captive marmosets housed at the Southwest National Primate Research Center (SNPRC), uncovering >90,000 SNPs^23^. We filtered these to identify 4,573 SNPs with high call rates and intermediate allele frequencies. We selected 560 regions spread over the autosomes of the marmoset genome to design a targeted amplicon sequencing platform using the rhAmpSeq system (SFig. 1A). The quality of this approach was high, in an initial sequencing run of 24 animals we detected a SNP genotype for 460/560 (82.1%) regions after stringent quality control. After further evaluation across 219 samples, we removed 26 SNPs with a minor allele frequency of <0.05. We retained 683 SNPs across 434 loci in the marmoset genome, with 156/434 (35.9%) of regions containing multiple SNPs. Depth of coverage across the retained SNPs was even (mean Gini coefficient of 0.3, SFig.1B). To estimate the accuracy of genotype calling, we looked at discordance across four tissues from a single individual with no evidence of chimerism. Here, we anticipate complete concordance between sequencing runs. Cumulatively, 4,059 of 4,077 (99.6%) genotypes matched, supporting the high accuracy of our platform.

We performed amplicon sequencing on 219 samples from a total of 82 animals from the SNPRC marmoset colony (Supplementary Table 1). These samples targeted 9 distinct tissues (hair follicle (HF, N=8), liver (LV, N=38), blood (BL, N=52), lymph node (LN, N=2), spleen (SP, N=5), adipose tissue (AD, N=29), kidney (KD, N=35), muscle (MS, N=39), skin (SK, N=8), and fecal samples (FC, N=3)). These tissues are predominantly derived from embryonic ectoderm (HF), embryonic endoderm (LV), lateral mesoderm (BL, LN, SP) and paraxial mesoderm (AD, KD, MS, SK), enabling the investigation of the developmental origins of chimeric material. We used amplicon sequencing for this analysis because it allowed us to exploit archived samples and pedigrees from our marmoset colony. For each sample we used the number of reads supporting the reference and alternate allele at each site to infer the within-sample allele frequency (WSAF) of each site. For each sample the WSAF is shown as a heatmap in Fig. 1A. For a non-chimeric sample, we expect that all heterozygous sites have a peak in WSAF around 0.5 on the y-axis. When samples contain a mixture of DNA from different sources, this peak can shift or flatten depending on the proportion of infiltrating DNA. We estimated the proportion of chimeric DNA using a maximum-likelihood method^24^. Fig. 1B-K shows the likelihood estimates for each sample across a range of chimeric fractions and divided by tissue. Each plot contains a curve for each sample showing the likelihood across the potential range of chimeric fraction (from no-chimerism to equal proportions of shared DNA). The peak in likelihood represents the most probable chimeric fraction value, a violin plot of this estimate is shown in Fig. 1L. These show that tissues derived from lateral mesoderm (predominantly from hematopoietic lineages) show the highest levels of chimerism, followed by paraxial mesoderm, embryonic endoderm and finally embryonic ectoderm. The data shown here provides a proportional estimate of the levels of chimerism present across different tissues and germ layers. The tissue-wide distribution of chimerism was previously explored in a closely related species (*Callithrix kulii*) by Ross *et al*.^12^ and Sweeney *et al*.^13,14^ (*C. jacchus*) who also found chimerism to be most prevalent in mesoderm derived tissues. SFig. 2 shows the agreement between Ross *et al*. and our data.

**Figure 1.**
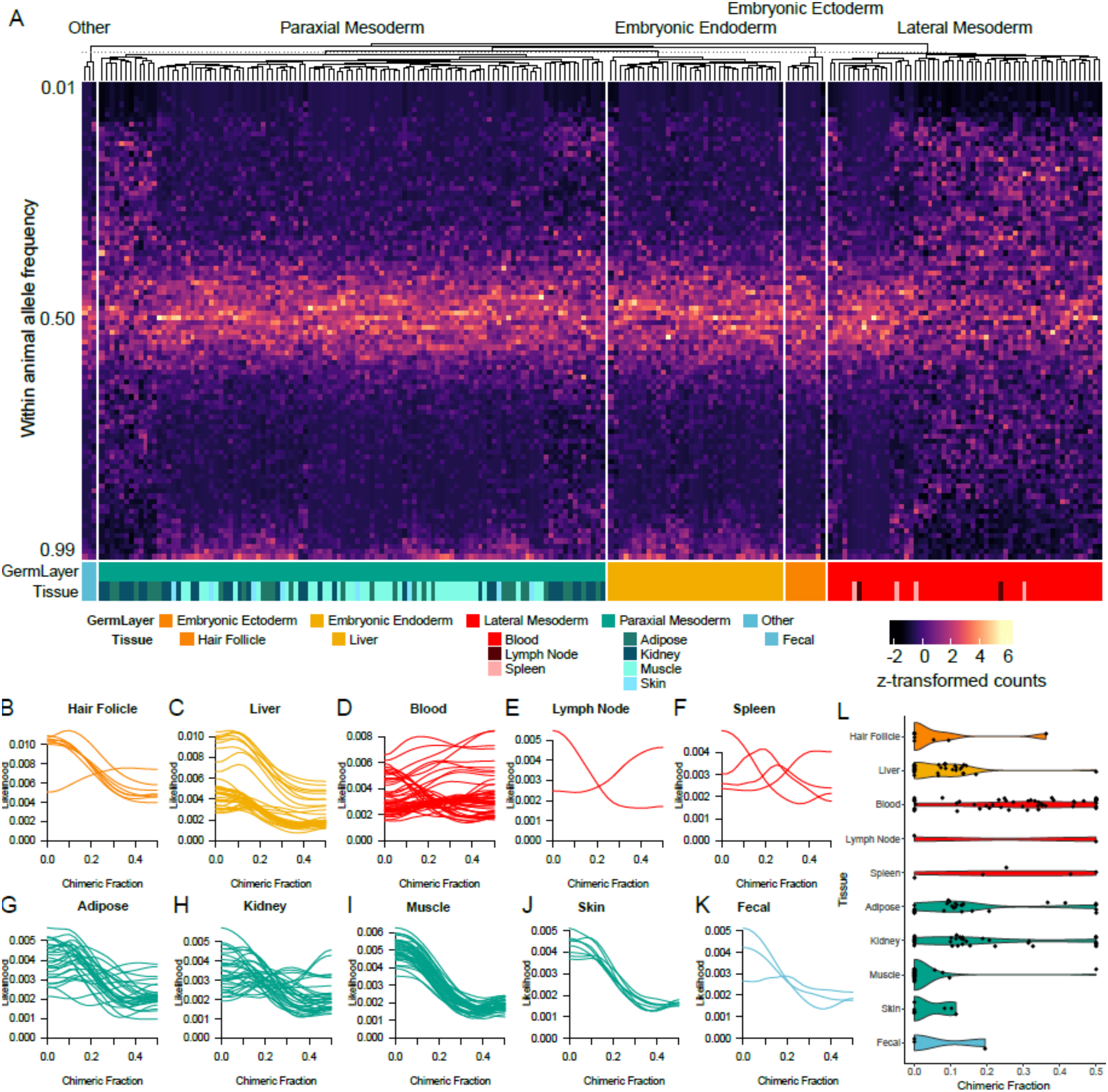
Assessing chimerism across 10 marmoset tissues. Upper panel. A heatmap showing the within-sample allele frequency of 683 SNPs called in our filtered dataset. Each row is a bin of allele frequency equaling 1%, and each column is the distribution of allele frequencies for a single sample. Within each major developmental lineage, samples are clustered, and color coded by both developmental lineage (Germ Layer) and terminal tissue (Tissue) sampled below the heatmap. (B-K) lower panel shows the likelihood estimates (y-axis) for the chimeric fraction (x-axis) within each sample with one curve per sample. (L) A violin plot of the chimeric fraction with the maximum likelihood for each sample.

**Figure 2.**
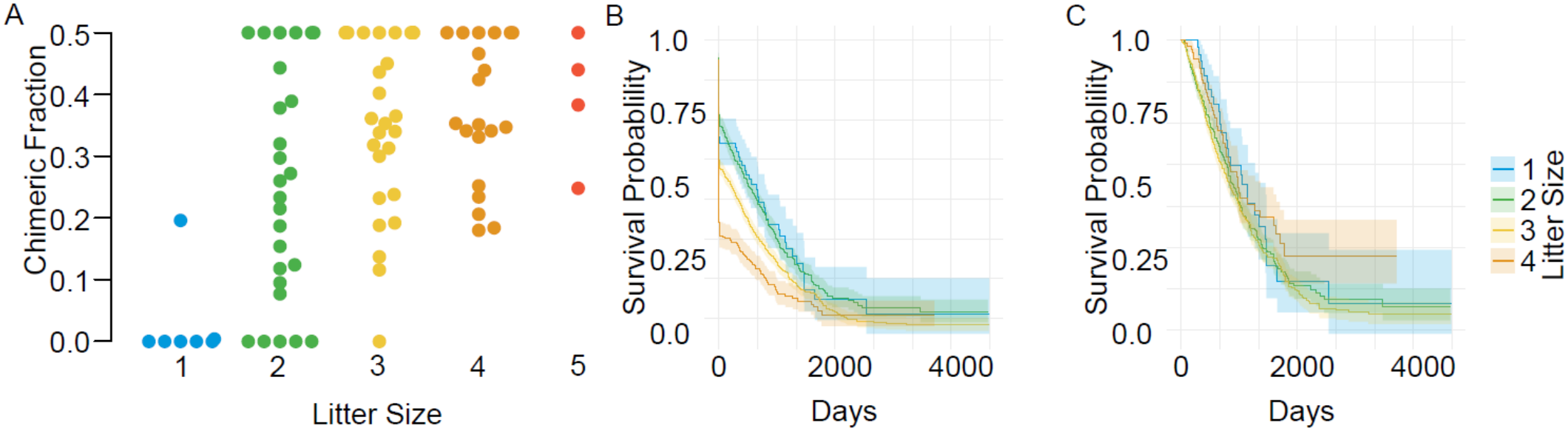
Chimeric fraction is associated with litter size. (A) A boxplot of chimeric fraction for each litter size at birth shows the chimeric fraction increases with litter size. (B) Survival curves for each litter size at birth shows a significant association between litter size and life span. (C) This effect is removed for animals surviving for more than 30 days. There were few animals born as quintuplets (n=30) and these have been removed from panels B and C for clarity. Full figures are in the supplementary materials.

### Litter size, but not age or sex predict chimeric fraction

To explore the relationship between chimeric fraction in the blood and age, sex or litter size at birth, we sequenced an additional 63 BL samples, creating a combined dataset of 115 estimates of BL chimerism. Increased litter size is significantly associated with higher chimeric fraction (linear regression, P=4.5×10^-5^, r=0.57). Neither age nor sex are associated with chimeric fraction. Our analysis of tissue-wide chimerism above included multiple tissues previously reported to show limited to no chimerism (such as hair follicles). Using these non-chimeric samples, we defined a threshold for chimerism as a chimeric fraction of ≥0.1. The analysis of litter size showed 1/7 (14.3%) of animals who were born without littermates had a chimeric fraction above this threshold in the blood. Given the high rate of multiples in marmosets^8^ this chimeric singleton is likely the result of absorption of littermates *in utero*, a finding supported by a previous study of marmoset prenatal sonography^25^. BL chimerism was observed in 27/28 (96.4%) of individuals born in a litter of 3 or more and 19/26 (73.1%) individuals of twin litters.

Our extensive pedigree records enable us to track the life span of animals from different litter sizes. There is a strong relationship between litter size and lifespan, with animals from higher litter sizes showing considerably reduced life spans (n=2,720, P<2.2×10^-16^, Cox Proportional Hazard Ratio=1.28). This is in close agreement with prior studies linking litter size to mortality in marmosets^26–28^. Notably, this effect appears limited to impacting survival during the first few weeks. After exclusion of animals who lived <30 days there is no impact of litter size on life span (n=1,246, p=0.5, Cox Proportional Hazard Ratio=0.98). As all our estimates of chimerism were obtained from animals surviving >25 days, we estimated the impact of chimerism on life span using a threshold of chimeric fraction>0.1. We did not see a significant impact (n=115, P=0.08, Cox Proportional Hazard Ratio=0.4), though the general trend towards non-chimeric animals surviving longer should be revisited with a larger sample size.

### Littermates show similar levels of chimerism in the blood

Examination of a single triplet litter by scRNAseq^29^ previously identified chimeric cells that were present from each littermate in all other littermates. We sought to determine whether this is always the case or whether chimeric cells are preferentially shared in one direction (Fig. 3A-B). Chimerism was present in every group of animals from the same litter we examined suggesting that the scenario in Fig. 3B is unlikely. To test this more quantitatively we first compared the allele frequencies in BL of full siblings from the same and different pregnancies to less related individuals (Fig. 3C) to see if littermates show more similar genetic composition than expected. Littermates had the most correlated BL allele frequencies (*r*^*2*^=0.91), followed by full-sibs (*r*^*2*^=0.51) and non-sibs (*r*^*2*^=0.15), with each comparison being significantly different. If littermates contain similar composition of sibling DNA we would expect them to show similar chimeric fractions. We compared the average difference in chimeric fraction (Δ chimeric fraction; Fig 3D) between littermates (0.097), full-sibs (0.15) and non-sibs (0.20), with littermates showing a significantly smaller difference in chimeric fraction compared to full-sibs (p=0.043, one-sided t-test) and non-sibs (p=0.002, one-sided t-test) and full-sibs significantly smaller than non-sibs (p=4.7x10-13, one-sided t-test).

**Figure 3.**
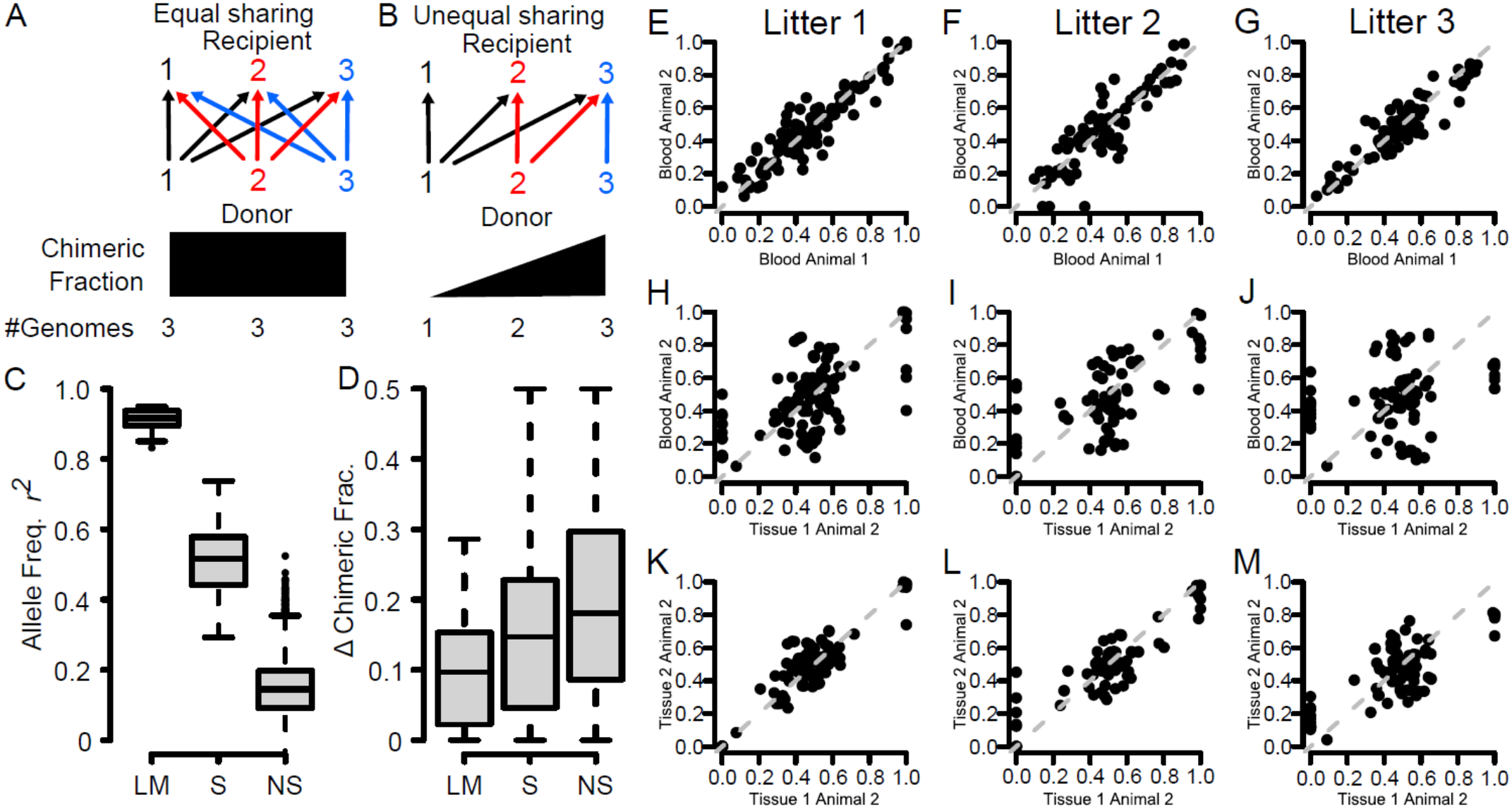
Sharing of chimeric cells between littermates. Two possible outcomes of cellular exchange *in utero*. In panel A all animals share equally in each direction, in panel B there is a strict hierarchy of sharing with animal 1 being a universal donor and animal 3 being a universal recipient. (C) A boxplot showing the correlation in allele frequency of blood between littermates (LM), non-littermate siblings (S) and non-siblings (NS). (D) A boxplot showing the difference in chimeric fraction of blood (ΔChimeric Frac.) between littermates, non-littermate siblings and non-siblings. (E-M) Scatter plots of unfixed allele frequencies within littermates comparing blood samples from different animals from the same litter (E-G), blood samples to tissue samples from the same animal (H-J) and tissue samples to tissue samples from the same animal (K-M). Each column shows data from the same litter.

While our examination of BL samples reveal they share a similar genetic composition within litters, it does not show if each individual from the litter is evenly represented. To explore this we sampled 5 pairs of littermates across blood and multiple tissues within minimal chimerism (Fig. 3E-M). For three litters we sampled blood from two littermates and matching tissue from one animal. This enabled us to compare the composition of the blood across littermates to the composition of tissue. In each case the blood samples were highly similar (*r*^*2*^=0.88-0.93), though the correlation between the allele frequencies in blood to tissue from the same animal was highly variable (Litter 1: mean *r*^*2*^=0.70 (0.65-0.73), Litter 2: mean *r*^*2*^=0.77 (0.67-0.93), Litter 3: mean *r*^*2*^=0.38 (0.19-0.66)). This highlights that the genetic composition of blood in an animal can be very distinct from the tissue and support that tissue genotypes can be poorly represented in the blood. For instance, in Litter 3 the composition of blood in animals 1 and 2 is highly similar (*r*^*2*^=0.93, Fig. 3G), though comparison of blood and tissue from the same animal show much less similar composition (*r*^*2*^=0.67, Fig. 3J).

Cumulatively, this supports multidirectional sharing of chimeric cells between littermates as in Fig. 3A.

### The genetic basis of chimerism and litter size

Prior work on identifying the genetics of litter size and chimerism in marmosets have focused on candidate genes or derived, fixed mutations along the Callitrichidae lineage^24,30,31^. In contrast, we find both litter size and chimerism to vary substantially in the SNPRC colony, suggesting there may be genetic variation segregating, which explains these traits. To begin to identify genetic architecture of litter size and chimerism we estimated the heritability of each trait. We distinguish here average litter size of a dam/sire (the average number of animals born in each litter to a dam or sire) from birth litter size (the litter size an animal is born into). We estimated the heritability of average litter size, birth litter size and chimerism using polygenic models in SOLAR using birth year, sex and litter size (for models testing chimerism) as covariates. For litter size estimates we used extensive pedigree and birthing records to substantially increase our sample size. All showed significant heritability (average litter size: n=508, *H*^*2*^=0.12, p=0.005; chimerism: n=112, *H*^*2*^=0.92, p=0.005; birth litter size: n=2,398, *H*^*2*^=0.55, p=8.3×10^-125^). We also ran sex specific estimates of heritability for average litter size which showed that females, but not males, have significant heritability (females: n=253, *H*^*2*^=0.19, p=0.026 – males: n=217, *H*^*2*^=0.03, p=0.38).

To refine our understanding of the genetic architecture of chimerism and litter size in marmosets, we performed whole exome sequencing of 379 marmosets from the SNPRC pedigree. Recent studies have explored the diversity and genetic structure of marmoset colonies showing moderate diversity and the presence of substantial differentiation between colonies^23,32,33^. To minimize the impact of chimerism on genotype inference, we used hair follicles as a source of DNA. We show here (Fig. 1) and previously^23^ that hair follicle DNA has minimal chimerism and can be captured non-invasively. We performed genome-wide association of chimeric fraction (as a binary trait with a threshold of 0.1), birth litter size and average litter size across available animals using GEMMA^34^. We plotted As with heritability, we performed association for average litter size both jointly and for each sex independently. At an FDR threshold of p<0.05, we identified four significant loci across three traits (Fig. 4). Quantile-quantile plots revealed minimal p-value inflation (SFig. 3). These include SNPs within the TMEM88B and ATAD3A genes for chimeric fraction, SNPs within the OPN3 gene for average litter size of dams and SNPs within the EPS8 and KCNQ3 genes for birth litter size. Further examination of the OPN3 locus showed clear segregation of average litter size by genotype (Fig. 4G) and showed that the reference allele of the most significant SNP has declined in frequency in the SNPRC colony over the last 20 years. We explored whether maternal OPN3 allele status correlated with chimeric fraction in offspring. We observed a trend towards higher chimeric fraction in the reference allele though the limited number of comparisons was likely not sufficient to reach significance.

**Figure 4.**
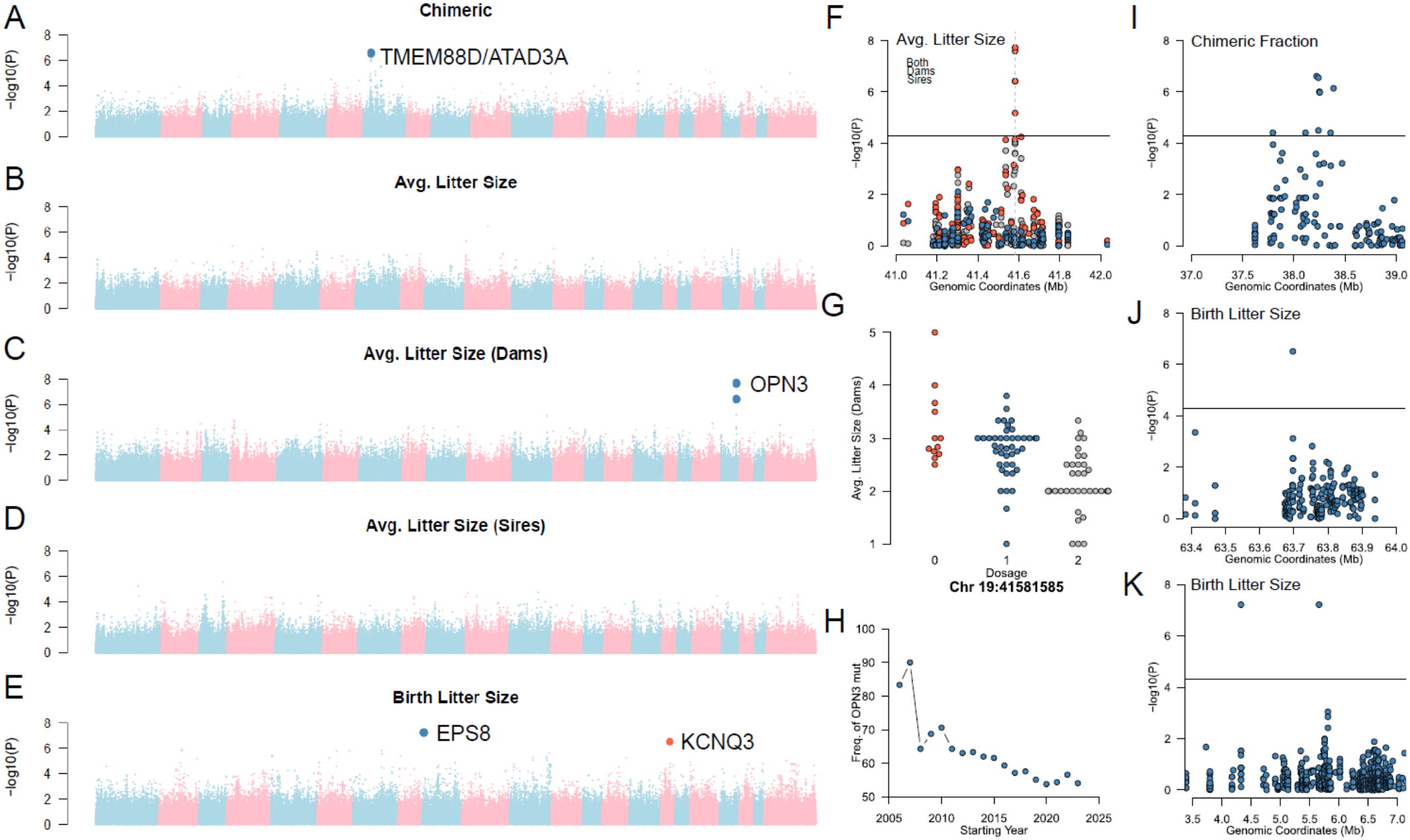
Genome-wide association of chimerism (A), average litter size (B) and average litter size of dams (C), average litter size of sires (D) and birth litter size (E). (F) The locus surrounding OPN3 for all parents (grey), dams (red) and sires (blue). (G) The average litter size of dams by allelic dosage at the most significant SNP (within the OPN3 gene). (H) The frequency of the non-reference allele changes over time in the sequencing data from the SNPRC colony. (I-K) The genomic context of the other hits (I; chimerism, surrounding TMEM88D/ATAD3A, J; birth litter size, surrounding EPS8 and K; birth litter size, surrounding KCNQ3).

## Discussion

We evaluated the genetics of chimerism and litter size in the marmoset, *Callithrix jacchus*. For decades, this trait has been a biological curiosity^35–38^ - however, the advent of highly sensitive sequencing approaches has identified chimerism to be widespread and clinically relevant in human health. For instance, fetal-maternal chimerism is now recognized as occurring widely across pregnancies^1,16,39^. Few well characterized models exist in which the genetics and outcome of chimerism can be evaluated. To aid in the development of the marmoset as a model of chimerism, we developed an accurate approach to quantify the fraction of cells from infiltrating littermates using amplicon sequencing within an existing maximum-likelihood framework. Cumulatively, single cell RNA-seq^29^, and PCR genotyping^12,13^ have suggested that chimerism is limited to the hematopoietic lineage. Using multiple tissues across 82 animals, our data is in broad agreement with these findings. We do not have the power with this study to deconvolute the tissues we sample into individual genotypes. However, we see that the chimeric fraction is correlated with the expected proportion of infiltrating hematopoietic lineage cells.

Our deep sampling of hematopoietically-derived lineages allows us to identify a relationship between the size of a litter an animal is born into and the degree of chimerism. As litter size increases, so does the likelihood of carrying cells from a littermate. As we see no link between age and chimeric fraction, these infiltrating sibling cells persist across the lifespan. The oldest animal in our dataset approaches the maximum recorded lifespan of a marmoset at 17 years and 10 months and exhibits chimerism, as do animals less than 2 years old. This is further supported by the comparison of littermates. We see remarkably similar genetic compositions of blood between littermates. Once chimerism is laid down during development, the proportion of cells appears to remain constant. The advent of spatial single cell RNAseq will be a rich avenue for mapping the direction and magnitude of chimerism in placental samples.

Our data enable the first glimpse into the genetic underpinnings of chimerism and litter size. Chimerism, birth litter size, and the average litter size of a dam all show significant heritability in our pedigree. We also identified significantly associated SNPs for each of these traits. Given the limited resolution and power afforded by the sample size and exome data we have used here, we cannot be certain about localizing the causal variants driving these traits. Scans for recent positive selection and long-term balancing selection can add support to association scans and aid in further refinement of causal variants. Two of the loci we identify here overlap with genes identified as the targets of positive selection in a recent study by Soni *et al*^40^. While we identify the opsin gene OPN3 as the lead candidate for average litter size of dams, the peak selected gene was RGS7. While either gene may play a role in modulating litter size, OPN3 is particularly intriguing. OPN3 perturbation in mice impacts litter size and neonatal development and has links to food intake and body temperature regulation^41,42^. Variation in OPN3 has also been associated with number of children and number of pregnancies in a cohort of Bangleshi women^43^. Similarly, we identify a region encompassing TMEM88B and ATAD3A as significantly associated with chimeric fraction. This overlaps with one of the most compelling signals from Soni *et al* - a peak surrounding the GNB1 gene.

The loci associated with birth litter size also contain interesting genes. While EPS8 and KCNQ3 do not have an obvious role in litter size, several of the flanking genes are compelling candidates for playing a role in pregnancy. At the EPS8 locus these include RERG, a regulator of estrogen levels^44^, ARHGDIB, a regulator of trophoblast invasion^45^, and MGP a modulator of smooth muscle and uterine function^46^. At the KCNQ3 locus these include a number of genes involved in regulating maternal-fetal interaction and trophoblast development including NDRG1^47,48^ and HHLA1^49^. Given the role of placentation in marmoset chimerism and patterns of prenatal growth in marmosets, the potential involvement of genes related to trophoblast function are intriguing^50,51^. While future work will be required to determine the specific genes and mechanisms involved these analyses provide compelling targets for further investigation.

## Methods

### Marmoset sampling

All animals were housed at the Southwest National Primate Research Center (SNPRC), Texas Biomedical Research Institute, an AAALAC accredited institution. Animals were maintained under standardized marmoset husbandry conditions^52^. The study was approved by the Texas Biomedical Research Institute Animal Care and Use Committee. The Ross lab maintains a marmoset frozen tissue bank of samples collected opportunistically at necropsy from colony clinical cases, flash frozen, and maintained at -80. Frozen whole blood samples are also maintained in the archive from collections during standard physical exams. Samples were selected from the archive for this project from individuals of known litter size, or from matched litter sets. All samples were blinded for initial genetic analyses. The SNPRC maintains all clinical health and demographic records in a Labkey based electronic health record. Additional blood samples from individuals of known litter size were collected for this project through the SNPRC biomaterials program.

### DNA Extraction

Extraction for all sample types was done with the QIACube and Qiagen Blood and Tussue kit (cat 69504). Standard QIACube protocols were followed for all tissues, and a bead beating step was added for adipose tissue to enhance extraction. DNA was eluted with 75uL of nuclease free water and measured by Qbit broad range reagents (Invitrogen).

### Amplicon Sequencing Protocol

DNA was diluted as necessary for aliquots of 0.9-9ng and run according to manufacturer’s directions (Integrated DNA Technologies - IDT) using their rhAMPSeq kit. Bead cleaning was performed after the Targeted PCR1 before the Indexing PCR 2 step. Samples were bead cleaned again and eluted in 22uL of IDTE, pH 8.0 (IDT). QC was performed by Agilent Tape Station D1000, and DNA was measured again by Qbit. Libraries were pooled based on Qbit values and the resulting pools were sent to Admera for sequencing on the NovaSeq platform.

### Alignment, variant calling, phasing and imputation

FASTQs were trimmed with fastp v0.22.0 along with the flags “-- detect_adapter_for_pe” and “--correction” specified. BAMs were then aligned to the Callithrix_jacchus_cj1700_1.1 reference genome with BWA-MEM v0.7.18-r1243-dirty and post-processed with SAMtools (samtools sort ${bam} -n -u | samtools fixmate - - -m -u | samtools sort - -o ${output}). The processed BAMs were then called for variants using GATK v4.6.1.0 tools. The first step was with HaplotypeCaller to create gVCFs, which in turn were converted into a GenomicsDB datastore with GenomicsDBImport, before being jointly called on autosomes with GenotypeGVCFs. Biallelic SNPs were kept from the resulting VCF with BCFtools v1.21 (bcftools norm ${vcf} -m-any --fasta-ref ${ref_fasta} -Ou | bcftools view -e’type!=“snp”‘ -Ou | bcftools norm -m+any -Ou | bcftools view -M2 -m2 -Ob -o ${output}). With BCFtools again, hard filtering was performed (bcftools filter ${input} -e ‘QUAL < 30 || QD < 2 || SOR > 3 || FS > 60 || MQ < 40 || MQRankSum < -12.5 || ReadPosRankSum < -8’ -Oz -o ${output}). GATK’s CalculateGenotypePosteriors and VariantFiltration were then used for genotype refinement. Further filtering with BCFtools removed any variants that became an AC of 0 from the refinement.

After variant calling we retained biallelic SNPs with <10% of missing data, and masked mendelian errors in plink v1.9^53^ (plink --bfile WES_${chr} --set-me-missing --mendel-duos --mendel-multigen --make-bed --out WES_ME_${chr} -- geno 0.1). We then phased genotypes using pedigree information using Shapeit2^54^ (shapeit -B WES_ME_${chr} -- duohmm -W 5 -O WES_${chr}_duohmm.phased) and imputed missing variants using Beagle v5.5^55^.

### Estimating the chimeric fraction

We estimated the chimeric fraction of each amplicon sequencing sample using a maximum-likelihood approach^24^ using a custom script. For each filtered biallelic SNP (DP>50, GQ>99) we estimated the within sample allele frequency as the proportion of reads carrying the non-reference allele.

### Survival Analysis

Survival analysis was performed using the Survival and ggsurvfit packages in R. Linear regression between age, sex and litter size was performed using the lm function in R. We extracted demographic information (birth year, age, sex, litter size at birth, offspring records for sires and dams) from labkey records at SNPRC and removed animals not born on campus. We calculated the average litter size as the mean of all litter sizes born to a parent.

Within- and between-sample allele frequency correlations were estimated using Pearson correlation.

### Heritability Analysis

We estimated broad-sense heritability in SOLAR^56^ using the polygenic function. For each trait we limited analysis to animals born at SNPRC and performed inorm normalization of traits and covariates.

### Genome-Wide Association

We estimated the relatedness matrix from standardized genotypes in GEMMA v0.94^34^. For chimeric fraction, we dichotomized the trait by setting a threshold of 0.1 for a sample being chimeric. We felt this better reflected the limitations of the ML approach. For birth litter size (the litter size an animal is born with) we excluded records for all animals not born at SNPRC. Using a linear model, we removed the effect of birth year and sex from litter size and used the residuals for GWAS. For average litter size (the average size of litters produced by a sire or dam), we excluded records for all animals not born at SNPRC, we then calculated the mean litter size for each dam and sire. We used a linear model to remove the effect of number of litters, birth year of sire/dam and sex and used residuals for GWAS.

## Data Availability

Amplicon sequencing data have been deposited in the Short Read Archive under accession code PRJNA1532492 Exome sequencing data have been deposited in the Short Read Archive under accession code PRJNA909825

## Ethics

All experiments with marmosets were performed in accordance with protocols approved by the Texas Biomedical Institutional Animal Care and Use Committee and adhere to the American Society of Primatologists Principles for the Ethical Treatment of Nonhuman Primates and complied with all applicable U.S. laws regarding animal research.

## Code Availability

Custom code for the maximum likelihood estimation of chimeric fraction analysis is available at https://github.com/cheesemania/MLE-of-Chimeric-Fraction

## Acknowledgements

We thank Dr. Tim Anderson and Dr. Suzette Tardif for their critical insight and staff at the Southwest National Primate Research Center.

