## Supplementary Material for "The genetic and environmental basis of hematopoietic chimerism in marmosets"

**Supplementary Figure. 1. Assessing the quality of the amplicon sequencing platform.** (A) the genome-wide distribution of regions included in the platform (blue) and SNPs typed within these regions (red). (B) the evenness of coverage across each sample. The dashed line represents where all amplicon regions would be sequenced to equal depth.

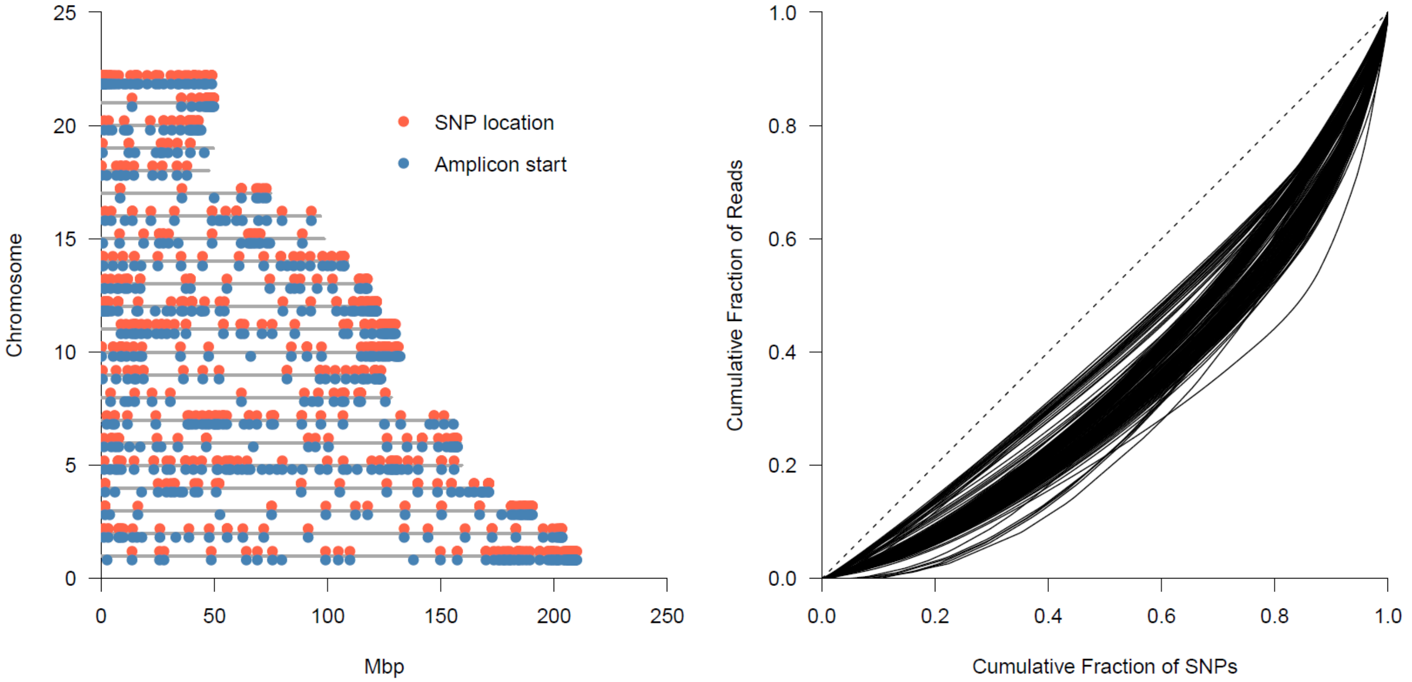

**Supplementary Figure 2. A comparison of the proportion of chimerism per tissue in Ross et al to this work.**

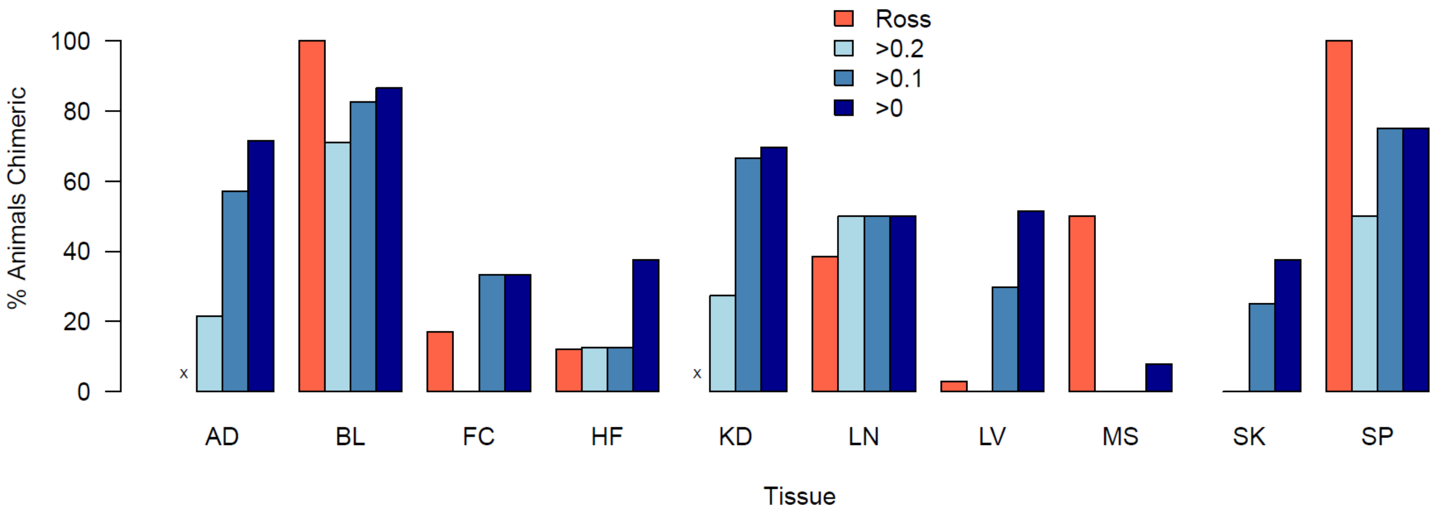

**Supplementary Figure 3. Quantile-quantile plots for genome-wide association of chimerism and litter size traits.** Genomic control values showed minimal inflation (Chimeric Fraction GC=1.49, Avg. Litter Size GC=1.03, Avg. Litter Size (Dams) GC=1.08, Avg. Litter Size (Sires) GC=1.04, Birth Litter Size GC=0.97).

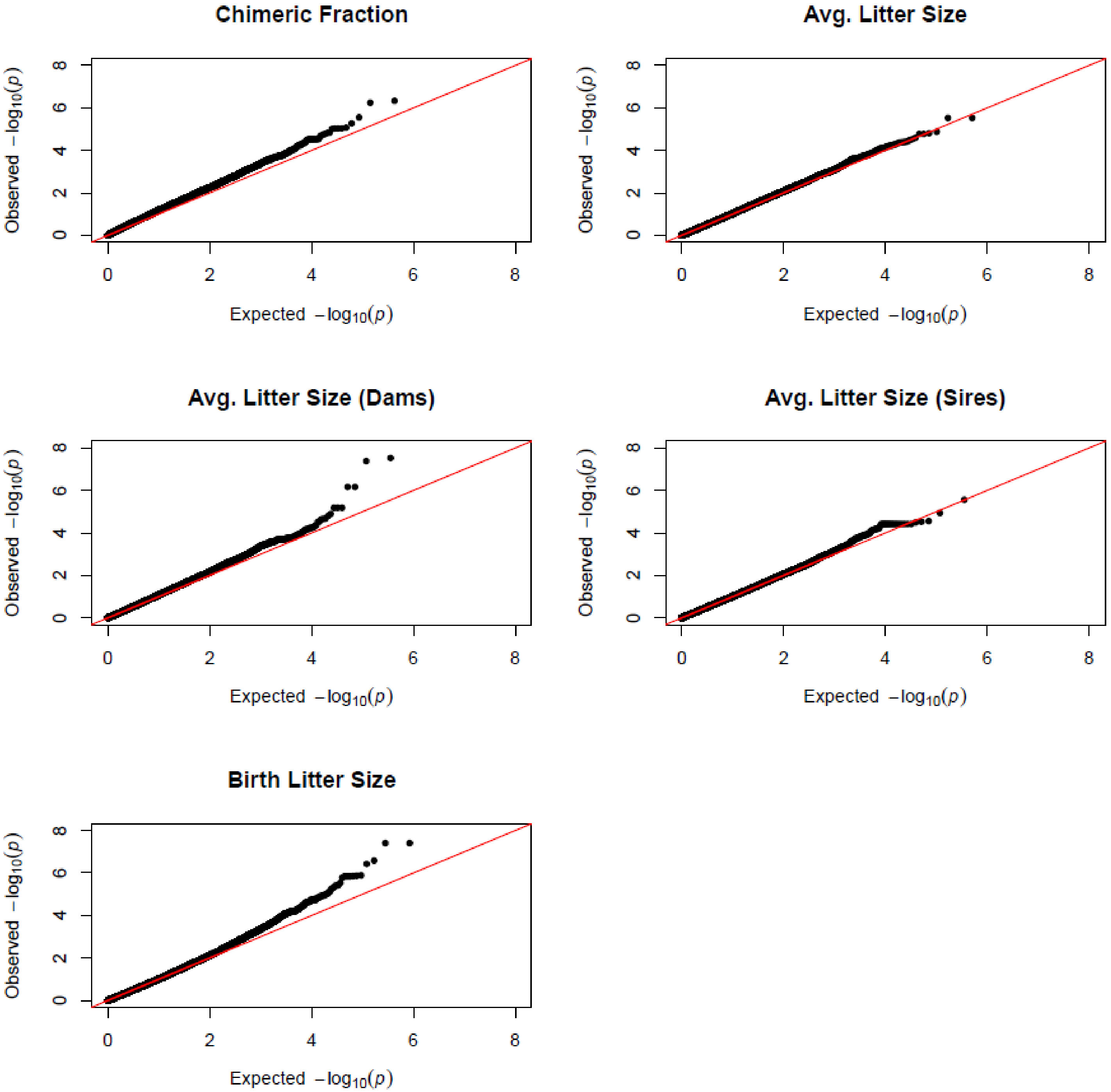

**Supplementary Table 1. Estimates of chimerism across tissue types**

| Chimeric Fraction | Tissue | AnimalID | Germ Layer | Age | Sex | Litter Size |
|---|---|---|---|---|---|---|
| 0.5 | LN | 1 | Lateral Mesoderm | 9 | M | 2 |
| 0 | MS | 1 | Paraxial Mesoderm | 9 | M | 2 |
| 0.43 | SP | 1 | Lateral Mesoderm | 9 | M | 2 |
| 0.5 | BL | 1 | Lateral Mesoderm | 9 | M | 2 |
| 0 | BL | 2 | Lateral Mesoderm | 8 | F | 1 |
| 0 | LN | 2 | Lateral Mesoderm | 8 | F | 1 |
| 0 | LV | 2 | Embryonic Endoderm | 8 | F | 1 |
| 0 | SP | 2 | Lateral Mesoderm | 8 | F | 1 |
| 0.113 | AD | 3 | Paraxial Mesoderm | 8 | F | 3 |
| 0 | LV | 3 | Embryonic Endoderm | 8 | F | 3 |
| 0 | MS | 3 | Paraxial Mesoderm | 8 | F | 3 |
| 0 | AD | 4 | Paraxial Mesoderm | 10 | F | 1 |
| 0.115 | LV | 4 | Embryonic Endoderm | 10 | F | 1 |
| 0 | MS | 4 | Paraxial Mesoderm | 10 | F | 1 |
| 0.117 | AD | 5 | Paraxial Mesoderm | 9 | M | 1 |
| 0 | MS | 5 | Paraxial Mesoderm | 9 | M | 1 |
| 0 | AD | 6 | Paraxial Mesoderm | 10 | F | 1 |
| 0 | LV | 6 | Embryonic Endoderm | 10 | F | 1 |
| 0 | MS | 6 | Paraxial Mesoderm | 10 | F | 1 |
| 0.201 | BL | 7 | Lateral Mesoderm | 17 | M | 2 |
| 0.253 | SP | 8 | Lateral Mesoderm | 13 | F | 1 |
| 0.097 | AD | 9 | Paraxial Mesoderm | 10 | M | 3 |
| 0.315 | KD | 9 | Paraxial Mesoderm | 10 | M | 3 |
| 0.131 | LV | 9 | Embryonic Endoderm | 10 | M | 3 |
| 0 | MS | 9 | Paraxial Mesoderm | 10 | M | 3 |
| 0 | SK | 9 | Paraxial Mesoderm | 10 | M | 3 |
| 0 | BL | 10 | Lateral Mesoderm | 13 | F | 2 |
| 0.253 | BL | 11 | Lateral Mesoderm | 11 | M | 2 |
| 0 | AD | 12 | Paraxial Mesoderm | 8 | F | 3 |
| 0.094 | HF | 12 | Embryonic Ectoderm | 8 | F | 3 |
| 0.116 | KD | 12 | Paraxial Mesoderm | 8 | F | 3 |
| 0.112 | LV | 12 | Embryonic Endoderm | 8 | F | 3 |
| 0.078 | MS | 12 | Paraxial Mesoderm | 8 | F | 3 |
| 0.238 | BL | 12 | Lateral Mesoderm | 8 | F | 3 |
| 0 | AD | 13 | Paraxial Mesoderm | 8 | F | 3 |
| 0 | HF | 13 | Embryonic Ectoderm | 8 | F | 3 |
| 0.135 | KD | 13 | Paraxial Mesoderm | 8 | F | 3 |
| 0.083 | LV | 13 | Embryonic Endoderm | 8 | F | 3 |
| 0 | MS | 13 | Paraxial Mesoderm | 8 | F | 3 |

| | | | | | | |
|---|---|---|---|---|---|---|
| 0.102 | SK | 13 | Paraxial Mesoderm | 8 | F | 3 |
| 0.361 | BL | 13 | Lateral Mesoderm | 8 | F | 3 |
| 0.5 | BL | 14 | Lateral Mesoderm | 10 | M | 3 |
| 0.5 | BL | 15 | Lateral Mesoderm | 11 | M | 3 |
| 0.116 | BL | 16 | Lateral Mesoderm | 10 | M | 3 |
| 0.5 | AD | 17 | Paraxial Mesoderm | 6 | F | 3 |
| 0.5 | KD | 17 | Paraxial Mesoderm | 6 | F | 3 |
| 0 | LV | 17 | Embryonic Endoderm | 6 | F | 3 |
| 0 | MS | 17 | Paraxial Mesoderm | 6 | F | 3 |
| 0.13 | AD | 18 | Paraxial Mesoderm | 6 | F | 3 |
| 0.314 | KD | 18 | Paraxial Mesoderm | 6 | F | 3 |
| 0.134 | LV | 18 | Embryonic Endoderm | 6 | F | 3 |
| 0 | MS | 18 | Paraxial Mesoderm | 6 | F | 3 |
| 0.339 | BL | 19 | Lateral Mesoderm | 10 | F | 3 |
| 0.09 | AD | 20 | Paraxial Mesoderm | 14 | M | 1 |
| 0 | KD | 20 | Paraxial Mesoderm | 14 | M | 1 |
| 0 | LV | 20 | Embryonic Endoderm | 14 | M | 1 |
| 0 | MS | 20 | Paraxial Mesoderm | 14 | M | 1 |
| 0 | BL | 21 | Lateral Mesoderm | 15 | F | 1 |
| 0.281 | BL | 22 | Lateral Mesoderm | 15 | F | 2 |
| 0.245 | BL | 23 | Lateral Mesoderm | 13 | M | 1 |
| 0.206 | AD | 24 | Paraxial Mesoderm | 9 | F | 1 |
| 0.326 | KD | 24 | Paraxial Mesoderm | 9 | F | 1 |
| 0 | LV | 24 | Embryonic Endoderm | 9 | F | 1 |
| 0 | MS | 24 | Paraxial Mesoderm | 9 | F | 1 |
| 0.5 | BL | 25 | Lateral Mesoderm | 12 | M | 1 |
| 0.33 | BL | 26 | Lateral Mesoderm | 12 | F | 2 |
| 0.107 | BL | 27 | Lateral Mesoderm | 11 | M | 2 |
| 0.267 | BL | 28 | Lateral Mesoderm | 11 | M | 3 |
| 0.235 | BL | 29 | Lateral Mesoderm | 11 | M | 2 |
| 0.081 | AD | 30 | Paraxial Mesoderm | 7 | M | 2 |
| 0.125 | KD | 30 | Paraxial Mesoderm | 7 | M | 2 |
| 0.17 | LV | 30 | Embryonic Endoderm | 7 | M | 2 |
| 0 | MS | 30 | Paraxial Mesoderm | 7 | M | 2 |
| 0.351 | BL | 30 | Lateral Mesoderm | 7 | M | 2 |
| 0.236 | BL | 31 | Lateral Mesoderm | 11 | F | 2 |
| 0.5 | BL | 32 | Lateral Mesoderm | 11 | M | 2 |
| 0.26 | BL | 33 | Lateral Mesoderm | 10 | F | 2 |
| 0.349 | BL | 34 | Lateral Mesoderm | 10 | F | 2 |
| 0.258 | BL | 35 | Lateral Mesoderm | 10 | F | 2 |
| 0.5 | BL | 36 | Lateral Mesoderm | 10 | F | 2 |
| 0.232 | BL | 37 | Lateral Mesoderm | 8 | F | 2 |
| 0 | AD | 38 | Paraxial Mesoderm | 19 | M | 1 |

| | | | | | | |
|---|---|---|---|---|---|---|
| 0.143 | KD | 38 | Paraxial Mesoderm | 19 | M | 1 |
| 0.085 | LV | 38 | Embryonic Endoderm | 19 | M | 1 |
| 0 | MS | 38 | Paraxial Mesoderm | 19 | M | 1 |
| 0.128 | AD | 39 | Paraxial Mesoderm | 18 | F | 1 |
| 0.222 | KD | 39 | Paraxial Mesoderm | 18 | F | 1 |
| 0 | LV | 39 | Embryonic Endoderm | 18 | F | 1 |
| 0 | MS | 39 | Paraxial Mesoderm | 18 | F | 1 |
| 0.104 | KD | 40 | Paraxial Mesoderm | 17 | M | 1 |
| 0 | LV | 40 | Embryonic Endoderm | 17 | M | 1 |
| 0 | MS | 40 | Paraxial Mesoderm | 17 | M | 1 |
| 0.135 | AD | 41 | Paraxial Mesoderm | 17 | F | 1 |
| 0.5 | KD | 41 | Paraxial Mesoderm | 17 | F | 1 |
| 0.135 | LV | 41 | Embryonic Endoderm | 17 | F | 1 |
| 0 | MS | 41 | Paraxial Mesoderm | 17 | F | 1 |
| 0.5 | KD | 42 | Paraxial Mesoderm | 12 | M | 1 |
| 0.5 | MS | 42 | Paraxial Mesoderm | 12 | M | 1 |
| 0.5 | SP | 42 | Lateral Mesoderm | 12 | M | 1 |
| 0 | KD | 43 | Paraxial Mesoderm | 11 | F | 1 |
| 0 | LV | 43 | Embryonic Endoderm | 11 | F | 1 |
| 0 | MS | 43 | Paraxial Mesoderm | 11 | F | 1 |
| 0.188 | SP | 43 | Lateral Mesoderm | 11 | F | 1 |
| 0.119 | KD | 44 | Paraxial Mesoderm | 16 | M | 1 |
| 0 | LV | 44 | Embryonic Endoderm | 16 | M | 1 |
| 0 | MS | 44 | Paraxial Mesoderm | 16 | M | 1 |
| 0 | BL | 45 | Lateral Mesoderm | 12 | M | 1 |
| 0.384 | BL | 46 | Lateral Mesoderm | 12 | M | 1 |
| 0 | BL | 47 | Lateral Mesoderm | 11 | M | 2 |
| 0.338 | BL | 48 | Lateral Mesoderm | 11 | F | 2 |
| 0.5 | BL | 49 | Lateral Mesoderm | 10 | M | 2 |
| 0 | BL | 50 | Lateral Mesoderm | 10 | M | 2 |
| 0.353 | BL | 51 | Lateral Mesoderm | 9 | F | 4 |
| 0.5 | BL | 52 | Lateral Mesoderm | 9 | F | 2 |
| 0.18 | BL | 53 | Lateral Mesoderm | 9 | F | 4 |
| 0.466 | BL | 54 | Lateral Mesoderm | 9 | F | 4 |
| 0 | KD | 55 | Paraxial Mesoderm | 5 | M | 2 |
| 0 | LV | 55 | Embryonic Endoderm | 5 | M | 2 |
| 0 | MS | 55 | Paraxial Mesoderm | 5 | M | 2 |
| 0.5 | BL | 56 | Lateral Mesoderm | 11 | M | 2 |
| 0 | BL | 57 | Lateral Mesoderm | 9 | M | 1 |
| 0 | BL | 58 | Lateral Mesoderm | 9 | M | 2 |
| 0.452 | BL | 59 | Lateral Mesoderm | 8 | M | 3 |

| | | | | | | |
|---|---|---|---|---|---|---|
| 0 | BL | 60 | Lateral Mesoderm | 8 | M | 3 |
| 0 | BL | 61 | Lateral Mesoderm | 8 | M | 2 |
| 0.341 | BL | 62 | Lateral Mesoderm | 8 | M | 4 |
| 0.367 | AD | 63 | Paraxial Mesoderm | 4 | M | 3 |
| 0 | HF | 63 | Embryonic Ectoderm | 4 | M | 3 |
| 0.132 | KD | 63 | Paraxial Mesoderm | 4 | M | 3 |
| 0.066 | LV | 63 | Embryonic Endoderm | 4 | M | 3 |
| 0.097 | MS | 63 | Paraxial Mesoderm | 4 | M | 3 |
| 0.5 | BL | 63 | Lateral Mesoderm | 4 | M | 3 |
| 0.5 | BL | 64 | Lateral Mesoderm | 8 | M | 3 |
| 0.5 | BL | 65 | Lateral Mesoderm | 8 | F | 4 |
| 0 | AD | 66 | Paraxial Mesoderm | 8 | F | 2 |
| 0 | KD | 66 | Paraxial Mesoderm | 8 | F | 2 |
| 0 | LV | 66 | Embryonic Endoderm | 8 | F | 2 |
| 0 | MS | 66 | Paraxial Mesoderm | 8 | F | 2 |
| 0.112 | AD | 67 | Paraxial Mesoderm | 8 | M | 2 |
| 0.188 | KD | 67 | Paraxial Mesoderm | 8 | M | 2 |
| 0 | LV | 67 | Embryonic Endoderm | 8 | M | 2 |
| 0.056 | MS | 67 | Paraxial Mesoderm | 8 | M | 2 |
| 0 | BL | 68 | Lateral Mesoderm | 8 | M | 2 |
| 0.362 | BL | 69 | Lateral Mesoderm | 8 | F | 1 |
| 0.206 | BL | 70 | Lateral Mesoderm | 8 | M | 4 |
| 0 | AD | 71 | Paraxial Mesoderm | 4 | M | 4 |
| 0.053 | HF | 71 | Embryonic Ectoderm | 4 | M | 4 |
| 0.114 | KD | 71 | Paraxial Mesoderm | 4 | M | 4 |
| 0.073 | LV | 71 | Embryonic Endoderm | 4 | M | 4 |
| 0.252 | BL | 71 | Lateral Mesoderm | 4 | M | 4 |
| 0 | BL | 72 | Lateral Mesoderm | 7 | M | 2 |
| 0.133 | AD | 73 | Paraxial Mesoderm | 4 | F | 4 |
| 0 | HF | 73 | Embryonic Ectoderm | 4 | F | 4 |
| 0 | KD | 73 | Paraxial Mesoderm | 4 | F | 4 |
| 0.067 | LV | 73 | Embryonic Endoderm | 4 | F | 4 |
| 0 | MS | 73 | Paraxial Mesoderm | 4 | F | 4 |
| 0.347 | BL | 73 | Lateral Mesoderm | 4 | F | 4 |
| 0 | BL | 74 | Lateral Mesoderm | 7 | F | 2 |
| 0.077 | BL | 74 | Lateral Mesoderm | 7 | F | 2 |
| 0.5 | AD | 75 | Paraxial Mesoderm | 4 | F | 2 |
| 0.206 | KD | 75 | Paraxial Mesoderm | 4 | F | 2 |
| 0 | LV | 75 | Embryonic Endoderm | 4 | F | 2 |
| 0 | MS | 75 | Paraxial Mesoderm | 4 | F | 2 |
| 0 | SK | 75 | Paraxial Mesoderm | 4 | F | 2 |
| 0.5 | BL | 76 | Lateral Mesoderm | 7 | F | 1 |

| | | | | | | |
|---|---|---|---|---|---|---|
| 0.5 | BL | 77 | Lateral Mesoderm | 7 | F | 2 |
| 0.5 | BL | 78 | Lateral Mesoderm | 7 | M | 2 |
| 0.5 | BL | 79 | Lateral Mesoderm | 6 | M | 3 |
| 0.5 | BL | 80 | Lateral Mesoderm | 5 | F | 2 |
| 0.5 | BL | 81 | Lateral Mesoderm | 7 | M | 4 |
| 0.5 | BL | 82 | Lateral Mesoderm | 7 | F | 4 |
| 0.5 | BL | 83 | Lateral Mesoderm | 7 | F | 4 |
| 0.5 | BL | 84 | Lateral Mesoderm | 7 | M | 3 |
| 0 | BL | 85 | Lateral Mesoderm | 6 | M | 2 |
| 0 | BL | 86 | Lateral Mesoderm | 6 | M | 3 |
| 0.378 | BL | 87 | Lateral Mesoderm | 6 | M | 2 |
| 0 | BL | 88 | Lateral Mesoderm | 5 | M | 1 |
| 0.297 | BL | 89 | Lateral Mesoderm | 5 | F | 2 |
| 0.198 | BL | 90 | Lateral Mesoderm | 6 | F | 1 |
| 0.5 | AD | 91 | Paraxial Mesoderm | 4 | M | 1 |
| 0.5 | KD | 91 | Paraxial Mesoderm | 4 | M | 1 |
| 0.5 | LV | 91 | Embryonic Endoderm | 4 | M | 1 |
| 0 | MS | 91 | Paraxial Mesoderm | 4 | M | 1 |
| 0.443 | BL | 92 | Lateral Mesoderm | 6 | M | 2 |
| 0 | BL | 93 | Lateral Mesoderm | 6 | F | 1 |
| 0.5 | BL | 94 | Lateral Mesoderm | 6 | M | 3 |
| 0.5 | BL | 95 | Lateral Mesoderm | 6 | M | 3 |
| 0.4 | BL | 96 | Lateral Mesoderm | 5 | F | 2 |
| 0.215 | BL | 97 | Lateral Mesoderm | 3 | F | 2 |
| 0.26 | BL | 98 | Lateral Mesoderm | 2 | F | 2 |
| 0.415 | AD | 98 | Paraxial Mesoderm | 2 | F | 2 |
| 0 | HF | 98 | Embryonic Ectoderm | 2 | F | 2 |
| 0.5 | KD | 98 | Paraxial Mesoderm | 2 | F | 2 |
| 0.086 | LV | 98 | Embryonic Endoderm | 2 | F | 2 |
| 0 | MS | 98 | Paraxial Mesoderm | 2 | F | 2 |
| 0 | BL | 99 | Lateral Mesoderm | 5 | M | 1 |
| 0.436 | BL | 100 | Lateral Mesoderm | 5 | F | 3 |
| 0.2 | BL | 101 | Lateral Mesoderm | 5 | M | 3 |
| 0.124 | BL | 102 | Lateral Mesoderm | 5 | M | 2 |
| 0.272 | BL | 103 | Lateral Mesoderm | 4 | M | 2 |
| 0.313 | BL | 104 | Lateral Mesoderm | 5 | F | 3 |
| 0.5 | BL | 105 | Lateral Mesoderm | 5 | F | 3 |
| 0.439 | BL | 106 | Lateral Mesoderm | 5 | M | 4 |
| 0.118 | BL | 107 | Lateral Mesoderm | 5 | M | 2 |
| 0.5 | BL | 108 | Lateral Mesoderm | 5 | F | 4 |
| 0.331 | BL | 109 | Lateral Mesoderm | 5 | F | 4 |
| 0.5 | BL | 110 | Lateral Mesoderm | 4 | M | 3 |
| 0.003 | BL | 111 | Lateral Mesoderm | 4 | F | 1 |
| 0 | AD | 112 | Paraxial Mesoderm | 1 | F | 1 |
| 0 | FC | 112 | Other | 1 | F | 1 |

| | | | | | | |
|---|---|---|---|---|---|---|
| 0 | KD | 112 | Paraxial Mesoderm | 1 | F | 1 |
| 0 | LV | 112 | Embryonic Endoderm | 1 | F | 1 |
| 0 | MS | 112 | Paraxial Mesoderm | 1 | F | 1 |
| 0 | SK | 112 | Paraxial Mesoderm | 1 | F | 1 |
| 0.103 | AD | 113 | Paraxial Mesoderm | 0 | F | 3 |
| 0 | KD | 113 | Paraxial Mesoderm | 0 | F | 3 |
| 0.136 | LV | 113 | Embryonic Endoderm | 0 | F | 3 |
| 0 | MS | 113 | Paraxial Mesoderm | 0 | F | 3 |
| 0.327 | BL | 114 | Lateral Mesoderm | 4 | M | 3 |
| 0.311 | BL | 115 | Lateral Mesoderm | 2 | M | 1 |
| 0.362 | HF | 115 | Embryonic Ectoderm | 2 | M | 1 |
| 0 | KD | 115 | Paraxial Mesoderm | 2 | M | 1 |
| 0.064 | LV | 115 | Embryonic Endoderm | 2 | M | 1 |
| 0 | MS | 115 | Paraxial Mesoderm | 2 | M | 1 |
| 0.5 | BL | 116 | Lateral Mesoderm | 4 | M | 5 |
| 0.248 | BL | 117 | Lateral Mesoderm | 3 | M | 5 |
| 0.34 | BL | 118 | Lateral Mesoderm | 4 | F | 3 |
| 0.32 | BL | 119 | Lateral Mesoderm | 4 | M | 2 |
| 0.159 | AD | 120 | Paraxial Mesoderm | 0 | F | 3 |
| 0.5 | KD | 120 | Paraxial Mesoderm | 0 | F | 3 |
| 0.126 | LV | 120 | Embryonic Endoderm | 0 | F | 3 |
| 0 | MS | 120 | Paraxial Mesoderm | 0 | F | 3 |
| 0.473 | BL | 121 | Lateral Mesoderm | 5 | F | 2 |
| 0 | KD | 122 | Paraxial Mesoderm | 0 | F | 3 |
| 0 | LV | 122 | Embryonic Endoderm | 0 | F | 3 |
| 0 | KD | 123 | Paraxial Mesoderm | 0 | F | 2 |
| 0.114 | LV | 123 | Embryonic Endoderm | 0 | F | 2 |
| 0 | MS | 123 | Paraxial Mesoderm | 0 | F | 2 |
| 0 | BL | 124 | Lateral Mesoderm | 4 | M | 1 |
| 0.5 | BL | 125 | Lateral Mesoderm | 3 | F | 3 |
| 0.5 | BL | 126 | Lateral Mesoderm | 17 | M | 1 |
| 0.5 | BL | 127 | Lateral Mesoderm | 12 | F | 1 |
| 0.5 | BL | 128 | Lateral Mesoderm | 10 | F | 1 |
| 0.093 | AD | 129 | Paraxial Mesoderm | 12 | M | 2 |
| 0.129 | KD | 129 | Paraxial Mesoderm | 12 | M | 2 |
| 0 | LV | 129 | Embryonic Endoderm | 12 | M | 2 |
| 0 | MS | 129 | Paraxial Mesoderm | 12 | M | 2 |
| 0 | SK | 129 | Paraxial Mesoderm | 12 | M | 2 |
| 0.165 | BL | 130 | Lateral Mesoderm | 12 | M | 2 |
| 0.108 | AD | 130 | Paraxial Mesoderm | 12 | M | 2 |
| 0.194 | FC | 130 | Other | 12 | M | 2 |

| | | | | | | |
|---|---|---|---|---|---|---|
| 0 | HF | 130 | Embryonic Ectoderm | 12 | M | 2 |
| 0.153 | KD | 130 | Paraxial Mesoderm | 12 | M | 2 |
| 0.084 | LV | 130 | Embryonic Endoderm | 12 | M | 2 |
| 0 | MS | 130 | Paraxial Mesoderm | 12 | M | 2 |
| 0 | MS | 130 | Paraxial Mesoderm | 12 | M | 2 |
| 0.083 | SK | 130 | Paraxial Mesoderm | 12 | M | 2 |
| 0.5 | AD | 131 | Paraxial Mesoderm | 11 | M | 2 |
| 0 | FC | 131 | Other | 11 | M | 2 |
| 0.099 | KD | 131 | Paraxial Mesoderm | 11 | M | 2 |
| 0.143 | LV | 131 | Embryonic Endoderm | 11 | M | 2 |
| 0 | MS | 131 | Paraxial Mesoderm | 11 | M | 2 |
| 0.114 | SK | 131 | Paraxial Mesoderm | 11 | M | 2 |
| 0.101 | KD | 132 | Paraxial Mesoderm | 10 | F | 1 |
| 0 | LV | 132 | Embryonic Endoderm | 10 | F | 1 |
| 0 | MS | 132 | Paraxial Mesoderm | 10 | F | 1 |
| 0.132 | KD | 133 | Paraxial Mesoderm | 11 | F | 1 |
| 0.105 | LV | 133 | Embryonic Endoderm | 11 | F | 1 |
| 0 | MS | 133 | Paraxial Mesoderm | 11 | F | 1 |
| 0 | SK | 133 | Paraxial Mesoderm | 11 | F | 1 |
| 0.341 | BL | 134 | Lateral Mesoderm | 3 | F | 4 |
| 0.5 | BL | 135 | Lateral Mesoderm | 3 | F | 4 |
| 0.44 | BL | 136 | Lateral Mesoderm | 3 | F | 5 |
| 0.383 | BL | 137 | Lateral Mesoderm | 3 | M | 5 |
| 0.425 | BL | 138 | Lateral Mesoderm | 3 | F | 4 |
| 0.5 | BL | 139 | Lateral Mesoderm | 3 | F | 2 |
| 0.5 | BL | 140 | Lateral Mesoderm | 3 | F | 3 |
| 0.365 | BL | 141 | Lateral Mesoderm | 3 | F | 3 |
| 0.409 | BL | 142 | Lateral Mesoderm | 2 | M | 2 |
| 0.354 | BL | 143 | Lateral Mesoderm | 2 | F | 4 |
| 0.352 | BL | 144 | Lateral Mesoderm | 2 | F | 4 |
| 0.5 | BL | 145 | Lateral Mesoderm | 1 | M | 4 |
